# Intra-exon motif correlations as a proxy measure for mean per-tile sequence quality data in RNA-Seq

**DOI:** 10.1101/2020.08.23.262055

**Authors:** Jamie J. Alnasir, Hugh P. Shanahan

## Abstract

Given the wide variability in the quality of NGS data submitted to public repositories, it is essential to identify methods that can perform quality control on these datasets when additional quality control data, such as mean tile data, is missing. This is particularly important because such datasets are routinely deposited in public archives that now store data at an unprecedented scale. In this paper, we show that correlating counts of reads corresponding to pairs of motifs separated over specific distances on individual exons corresponds to mean tile data in the datasets we analysed, and can therefore be used when mean tile data is not available.

As test datasets we use the *H. sapiens* IVT (*in-vitro* transcribed) dataset of Lahens et al., and a *D. melanogaster* dataset comprising wild and mutant types from Aerts et al.

The *intra-exon* motif correlations as a function of both GC content parameters are much higher in the *IVT-Plasmids* mRNA *selection free* RNA-Seq sample (control) than in the other RNA-Seq samples that did undergo mRNA selection: both ribosomal depletion (*IVT-Only*) and PolyA selection (*IVT-polyA*, wild-type, and mutant). There is considerable degradation of similar correlations in the mutant samples from the *D. melanogaster* dataset. This matches with the available mean tile data that has been gathered for these datasets. We observe that extremely low correlations are indicative of bias of technical origin, such as flowcell errors.

## 1 Introduction

Next-generation sequencing (NGS) methods have revolutionised nucleic acid sequencing largely as a result of the employment of fluorescence-based nucleotide chemistry to generate a light signal on nucleotide incorporation [1, 2, 3], miniaturisation and massively-parallel sequencing reactions [4]. Though these have, to a degree, simplified the core sequencing process allowing reactions to be performed in clusters to generate enough signal and in parallel to increase throughput, NGS technologies share the same complex preparatory procedures [5]. These are typically i) fragmentation of fragments to the size appropriate for the target sequencing platform, amplification (e.g. PCR), and ligation of synthetic sequencing adapters for the sequencing platform. Such high-throughput sequencing technologies generate millions to billions of reads in a matter of days and generate large datasets [6] — it has enabled a number of large-scale sequencing projects. These include the 100,000 genomes project in the UK [7, 8], the NIH (National Institute of Health) Precision Medicine 1 million genomes project in the US [9], and a 1 million genome project by the BGI (Beijing Genome Institute) in China [10] to name just a few. Global collaboration, such as that of the International Cancer Genome Consortium (ICGC), which coordinates cancer genomics research across different nations [11] is now possible thanks to the availability of Next-generation sequencing technology.

RNA-Seq, on which this work focuses, is a high-throughput NGS technique for estimating the concentration of *all* transcripts in a transcriptome. This is in contrast to microarrays, which are constrained to identification and quantification of pre-selected target sequences based on complementary probes immobilised on the array [12]. It provides wider coverage of the transcriptome as its methods involve the direct sequencing of transcripts of RNA found in the sample [13, 14]. RNA-Seq can, therefore, be used to study various types of RNA present: total RNA, mRNA, pre-mRNA, and non-coding RNA (ncRNA), such as microRNA and long ncRNA enabling it to be used to study alternative splicing events [15, 16]. Furthermore, RNA-Seq achieves this at a higher resolution [14] than other technologies. After applying RNA-Seq, the transcriptome can then be constructed by *mapping* read data back to a reference genome (a process involving the *alignment* of sequences in the read data to the reference). To quantify gene expression, this mapping process is combined with gene boundary information so as to count the number of transcripts that map to a given gene or exon region [17, 13, 18].

RNA-Seq has transformed our view of the extent and complexity of the transcriptome through deep-sequencing [13] and also as a result of the increased precision the technique offers over other methods. Whilst recent developments in the RNA-Seq workflow, from sample preparation to sequencing, have furthered our understanding of the transcriptome, they have also required substantial effort for data analysis and computation, and given the complexity of RNA-Seq workflow necessitates study of the bias that can be introduced in the preparatory steps [14, 19]. Characterisation of bias in RNA-Seq is especially incumbent given that the method sequences and measures the transcriptome indirectly using reverse-transcribed complementary DNA (cDNA) [20]. Whilst microarray technologies also suffer from unwanted sources of bias — in particular, the hybridisation of probes is known to lack specificity leading to increased variability — these sources are well characterised and estimates of gene expression have, therefore, been amenable to improvements by the application of statistical techniques. RNA-Seq technology, however, is comparatively younger and sources of bias and variability are still under investigation. Bias introduced in the preparatory steps can have a profound effect on the raw data and typically manifest themselves as sequence-specific or positional biases, whilst bias introduced by the sequencing process itself are often systematic in nature [21].

The main obstacle to obtaining accurate estimates of transcript expression from RNA-Seq data is non-uniformity in the distribution of mapped reads to the reference genome, which reduces the certainty that the measured counts of mapped reads reflect the true expression of the transcript within the cell’s transcriptome. These bias have numerous sources such as, for example, *wet-lab* sample preparatory techniques, the sequencing process itself [22, 23] and the potential for errors in post-sequencing data processing. They perturb the uniformity of the distribution of mapped reads to a reference genome [24] and such bias manifests itself as sequence-specific or positional [25]. Also, positional biases can occur due to random hexamer priming in sample preparation [26]. Large amounts of this raw RNA-Seq read data is deposited in public repositories such as the Sequence Read Archive (SRA) [27] and Gene Expression Omnibus (GEO) [28]. Furthermore, the SRA, for example, does not require a quality check on submission [29], and has shown poor annotation of sequencing protocol steps — both at the top-level study and individual experiment record level [23]. Hence it is critical that methods are developed to characterise and quantify bias in these datasets. Such methods can augment the analysis of QC metadata in the datasets or can serve as an alternative measure when this metadata is not present.[30] In this paper, we propose that a method we have previously devised, which applies distributed-computing to quantify the sequence-specific deviations in the uniformity of mapped reads [31], can be used as a proxy measure when mean tile data is not available for short-read RNA-Seq data. Our method uses counts of reads overlapping motifs and works at the deep, read level. This approach is based on the assumption that *4-mers* in short reads from one region on an exon will be correlated with *4-mers* in short reads from another region of the same exon. In order to provide the capacity to process the amounts of data typical in transcriptomic datasets [32] our analysis employs parallel distributed computing algorithms and infrastructure using the Apache Spark platform — it is named Hercules [33] (an MPI version has now also been made available). We demonstrate this using a controlled *in-vitro transcribed* (IVT) dataset created by Lahens et al. for the purpose of characterising bias introduced in RNA-Seq library preparation [24], as well as a *D. melanogaster* dataset.

The first dataset (IVT) was produced utilising *in-vitro transcription* in *E. coli* to clone a pool of approximately 1,000 pre-selected human plasmids from the Mammalian Gene Collection (MGC) [34]. Because the sequences and expression levels of these plasmids are known, and they do not undergo splicing, this allowed them to generate a highly controlled set of samples, and therefore a controlled dataset, in which the source of biological variation in the samples is minimised. The samples in this dataset were then subjected to different RNA-Seq preparatory protocols, specifically varying the step in which mRNA is selected. This enables the study and quantification of the effect of these steps on coverage levels of the MGC transcripts when they were aligned to the human reference genome (hg19/grch37). They found that the mRNA selection methods employed in RNA-Seq protocols, poly-A and ribosomal depletion, both resulted in significant fold changes in the coverage of the IVT MGC plasmids when compared to sequencing the IVT MGC plasmids directly (without mRNA selection). Importantly, as the bias introduced in this dataset is well characterised and attributed, we apply our analysis method to a selection of relevant samples.

In previous work, we applied our analysis method to replicates of two samples from a *D. melanogaster* dataset produced from typical biological specimens using conventional RNA-Seq protocols [31]. These are two small, but whole transcriptomes — those of the fruit fly species *D. melanogaster* wild-type and mutant-r2, comprising of approximately 12.9 M and 15.0 M reads respectively. We will re-examine our analysis of this data with respect to sequencing tile means. Whilst the IVT dataset offers a set of samples that allow us to study the effect of different RNA-selection methods, the *D. melanogaster* datasets have the same RNA-selection method applied to all samples and vary only in the glass eye mutation — i.e. the technical variation is fixed, and the biological variation should be minimal. Furthermore, the Drosophila species and its reference genome are extremely well studied and annotated, and the data has excellent provenance.

## 2 Materials and methods

### 2.1 Quantifying sequence-specific deviation in the distribution of mapped reads across exons

The uniformity of read distribution across an exon can be quantified by computing Pearson (or Spearman rank) correlations of the counts for the given motif pair in all exons within the dataset by aggregating the motif pair counts at a given distance apart (motif-spacing) regardless of position within the exon. We used motif-spacings of 10, 50, 100 and 200 base-pairs (bp). An ideal dataset would have perfect correlations for motif pairs (for instance +1.0 for the Pearson correlation coefficient) for any given motif-pair and motif-spacing. In order to thoroughly examine the affect of sequence-specific motifs on the uniformity of read distribution we analysed the correlation for all *4-mer* motifs ranging from AAAA to GGGG (i.e. 4^4^ combinations) in the RNA-Seq reads of an aligned SAM (Sequence Alignment Map) file. We verified this method by running our analysis on an artificially created transcriptome with in-built uniform distribution of reads (see Supplementary Information Section 1 and Figure S1).

The effect of extremes of GC content in RNA-Seq data (as well as microarray data) has been discussed in numerous studies [35, 36], and we therefore also investigate the effect the mean GC content of reads within the exon 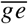, and the GC content of the *4-mer* motif itself *gm*, has on the distribution of reads across the exon. In order to partition reads by mean GC content we define binned GC content ranges: 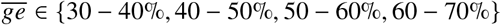. Given we are working with *4-mers*, the motif GC is a value in the set: *gm* ∈ {0%, 25%, 50%, 75%, 100%}.

### 2.2 *H. sapiens* IVT (*In-vitro* Transcription) RNA-Seq dataset

We explored known-bias in this RNA-Seq, by analysing *intra-exon* motif pair correlation within the reads, we performed analysis on three samples from *H. sapiens* that were produced in a controlled way using IVT (in vitro transcription), and by applying different library preparation protocols to each sample during RNA-Seq. These samples from Lahens et al. are known to demonstrate *intra-exon* coverage bias (we have used their *H. sapiens* data) [24]. Although the raw data deposited in GEO for this dataset has not been aligned, the library preparation protocol for each sample, and the alignment and post-processing strategy applied have been clearly documented. We, therefore, aligned the reads of the samples to the reference *hg19* genome according to the documented parameters to generate the SAM files for analysis by our method.

The RNA in these samples has been transcribed from cDNA clones in *E. coli* DH5α cells. The dataset comprises of a pool of 1,062 RNAs from a full-length human cDNA library sequenced using RNA-Seq. The first sample, *IVT-Only*, had its IVT RNA subjected to ribosomal RNA depletion prior to sequencing, whilst the second sample, *IVT-PolyAsel*, had polyadenylated selection applied instead of ribosomal depletion — these are two different, routinely used protocols for selecting specifically mature (mRNA) from RNA samples. The third sample, *IVT-Plasmids*, is our *control* as it was produced by direct sequencing of the Human IVT plasmids without RNA-selection (i.e. neither ribosomal depletion nor polyA selection were applied). The datasets were produced by the Smilow Center for Translational Research, Philadelphia, USA and featured in a publication by Lahens et al. [24], and the third sample was one of the controls used in the original research paper. This data is deposited at the GEO database with the ID GSE50445 [37].

The distribution of protein-coding exon lengths in *H. sapiens* is shown in Supplementary Figure S2. We note that, although the median exon length in *H. sapiens* is 121 bp (shorter than Drosophila) and approximately 80% of the exons are less than 200 bp (Sakharkar et al., 2004), the remaining 20% of exons will contribute to motif pair correlations at 200 bp apart.

### 2.3 *D. melanogaster* RNA-Seq dataset

In order to investigate *intra-exon* motif pair correlation within the reads from a “typical dataset”, we have used two Drosophila (species *D. melanogaster*) transcriptomics datasets which differ by mutation *gl[60j]* in the eye-antennal disc. These are the full transcriptomes of the wild-type and mutant glass eye mutations, acquired from Stein Aerts Laboratory of Computational Biology at the University of Leuven, Belgium. This data is deposited at the GEO database with the ID GSE39781 [38]. The *D. melanogaster* datasets featured in a research publication by Naval-Sánchez et al.[39].

Supplementary Figure S3 shows the distribution of exon lengths in the Drosophila genome — the median exon length is 298 bp. This is important because it shows that most of the exons are longer than 200 bp and therefore can have data to compute correlations.

The information regarding the source of the biological samples, the sample preparatory protocols and post-sequencing processing that were applied to samples in these two datasets are documented in Table 1.

**Table 1:** The sample and library preparation protocols, together with the data-processing steps, applied to the RNA-Seq datasets used in this analysis. (Left) *H. sapiens* IVT RNA-Seq. (Right) *D. melanogaster*.

| *H. sapiens* (IVT-Plasmids, IVT-Only and IVTpolyAseI) | *D. melanogaster* (wild and mutant) |
| --- | --- |
| **Replicates:** 1x .sam file for each IVT sample (no replicates) | **Replicates:** 4x .sam files (two replicates for each species). |
| **Sample preparation** | **Sample Preparation** |
| Glycerol stocks containing individual cDNAs (cloned into pCMV-Sport 6 plasmid) from the MGC (Mammalian Gene Collection) [34] were produced. Plasmid DNA was extracted from these glycerol stocks and plated at 50 ng per well in 384-well plates. The contents of three 384-well plates (total of 1,062 human transcripts) were collected. The plasmid library was then amplified by transferring 10 ng into *E. coli* DH5α cells (Invitrogen, Life Technologies, Carlsbad, CA, USA, catalogue no. 18258-012). The heat shock method was used to transform *E. coli* (See [24] for more details). The plasmids were then purified using Qiagen (Hilden, Germany) maxiprep kit (catalogue no. 12163), according to the manufacturer’s protocol. Samples were sequenced on the Illumina HiSeq 2000 platform. | For the wild type *D. melanogaster*, fly stocks (Canton-S and strain RAL-208) were obtained from the inbred collection of T. Mackay. For the mutant-r2 *D. melanogaster*, fly stocks (stock 507) were obtained from the Bloomington Stock Center. Eye-antennal and wing imaginal discs were dissected. RNA was extracted, yielding ~ 3 mg of total RNA per sample. The samples were processed into libraries according to the Illumina TruSeq protocol with appropriate indices, pooled. Sequencing of the transcriptomes was performed on the Illumina HiSeq 2000 platform. |
| **RNA-Seq** | **RNA-Seq** |
| After sequencing, raw reads from the samples were aligned to the human genome (GRCh37/hg19) using the RNA-Seq Unified Mapper (v2.0.4) with default parameters. Only reads that mapped to a single location were used (selected from the *RUM_Unique* aligned reads file). | After sequencing, Fastx-clipper [40] was used to discard reads containing residuals of adapter sequences were discarded (FastX clipper version 0.0.13 with option -M15). Quality control was applied to the raw sequence reads and performed using the FastQC software [30] (version 0.9), checking for PHRED quality >20 and different primer contaminations. The reads were then aligned using TopHat v2.0 [41] with default parameters, to the Flybase Drosophila Melanogaster genome version r5.45 (released March 2012) [42]. |

## 3 Results

### 3.1 Analysis of IVT (*In-Vitro* Transcribed) RNA in *H. sapiens*

We have analysed three *H. sapiens* samples which were prepared by IVT RNA-Seq. Lahens et al. in their study used RNA that has been in-vitro transcribed (IVT) from cDNA clones in *E. coli*[24]. Their rationale was that the “nucleotide sequence at every base was known, the splicing pattern established, and the expression the level coverage is uniform across the transcript.”. This means that any bias occurring in the coverage of reads in these three samples must be as a result of technical rather than biological origin.

The first IVT sample we analysed was the *IVT-Plasmids* sample, as this was produced from sequencing the Human IVT plasmids directly without applying Ribosomal depletion or PolyA selection methods, and therefore represents a control. Importantly, the *IVT-Plasmids* sample, by virtue of not having RNA selection protocol steps applied, also reduces the technical sources of variation in read distribution. Table 2 shows a number of *4-mer* motif-pairs that have very high correlations (Pearson correlations very close to +1), and these high correlations are observed across all spacings. There are also some extremely low correlations due to a lack of *4-mer* data (as indicated by the sample sizes in parenthesis). In order to visualise correlations across the *IVT-Plasmids* sample, we partitioned the results as a function of GC content of the motif and GC content of the exon and produced box and whisker plots (Figure 1). We observe a reasonably good correlation, with a median correlation of approximately 0.35, across all of the *4-mer* pairs as a function of the GC content of the exons (right side of the figure), and a reasonably good correlation across different Motif GC concentrations except that of 100% motif GC content.

**Table 2:**
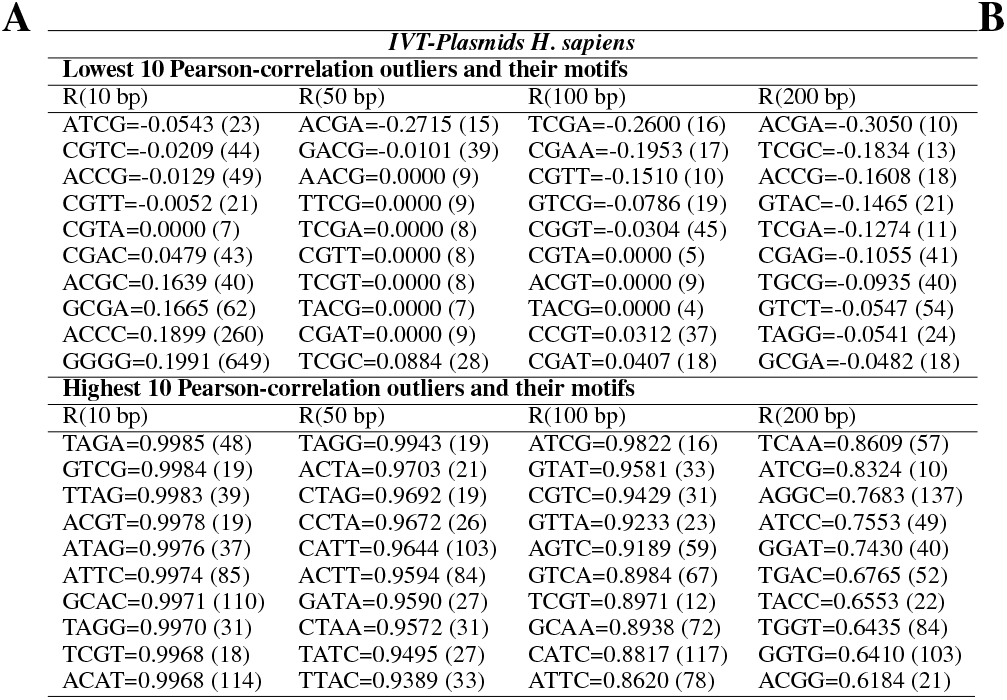
*H. sapiens* Pearson correlation co-efficient outliers (top ten and lowest ten) for different *intra-exon 4-mer* motif sequence pairs at 10, 50, 100 and 200 bp spacings.

**Figure 1:**
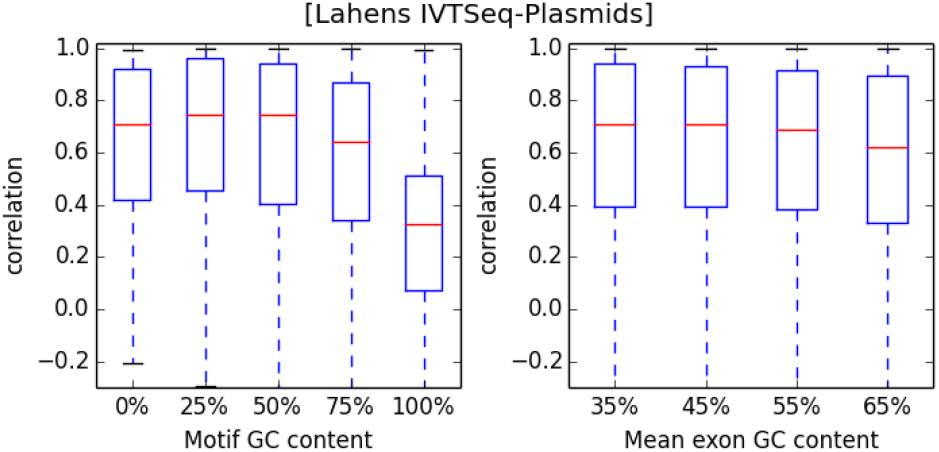
Box-whisker plot of the *IVT-Plasmid* sample (control) correlations as a function of Motif GC and mean exon GC content.

In order to compare the effect of applying different RNA selection protocol methods to the IVT samples, specifically ribosomal depletion vs. polyA selection, we compared the *IVT-Only* and *IVT-PolyA* samples respectively. Table 3 below shows that the highest outliers for both these IVT-Seq samples show a number of *4-mer* motif-pairs that have very high correlations. High correlation outliers are observed across all spacings. We produced box-whisker plots of correlation as a function of GC content of the motif and GC content of the exon (Figure 2). The trend of the data for *IVT-Only* and *IVT-PolyA*, as a function of Motif GC content and Mean GC content, is similar to that of the *IVT-Plasmids* data although the median correlation is somewhat less.

**Table 3:**
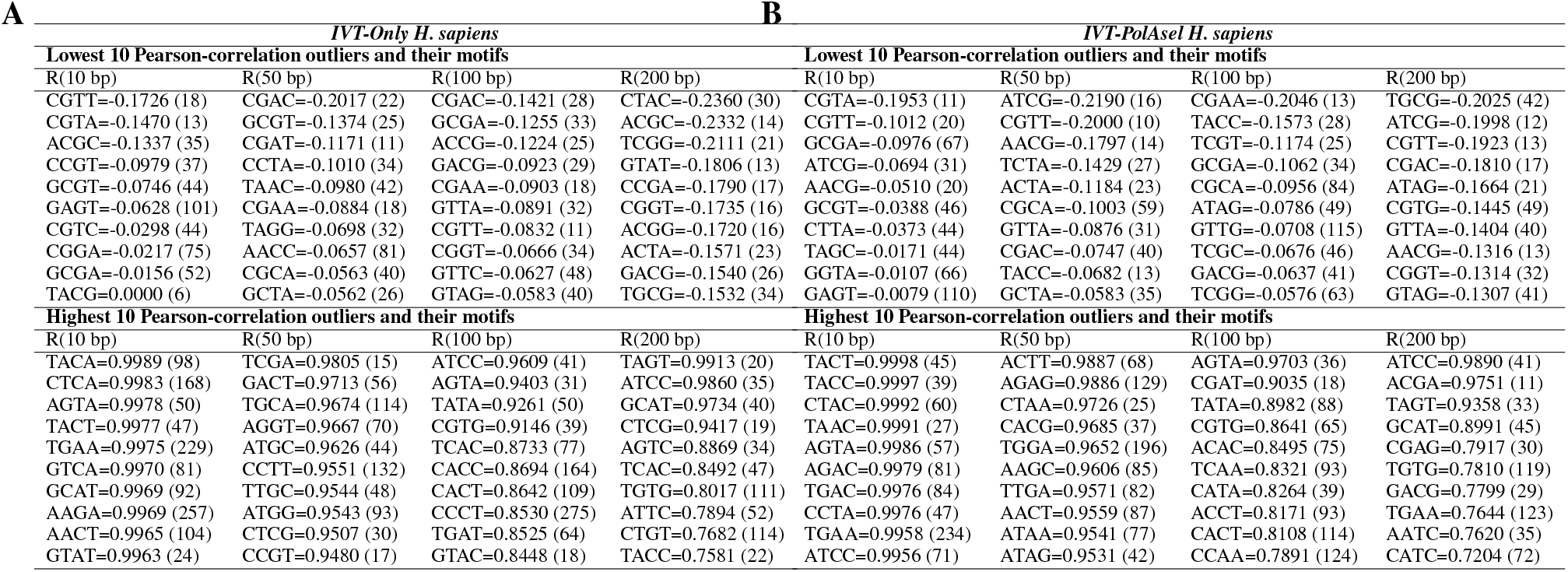
*H. sapiens IVT-Seq* Pearson correlation co-efficient outliers (top ten and lowest ten) for different *intra-exon 4-mer* motif sequence pairs at 10, 50, 100 and 200 bp spacings. A) IVT only library preparation B) IVT with PolyA library selection. Sample sizes are given in parenthesis.

**Figure 2:**
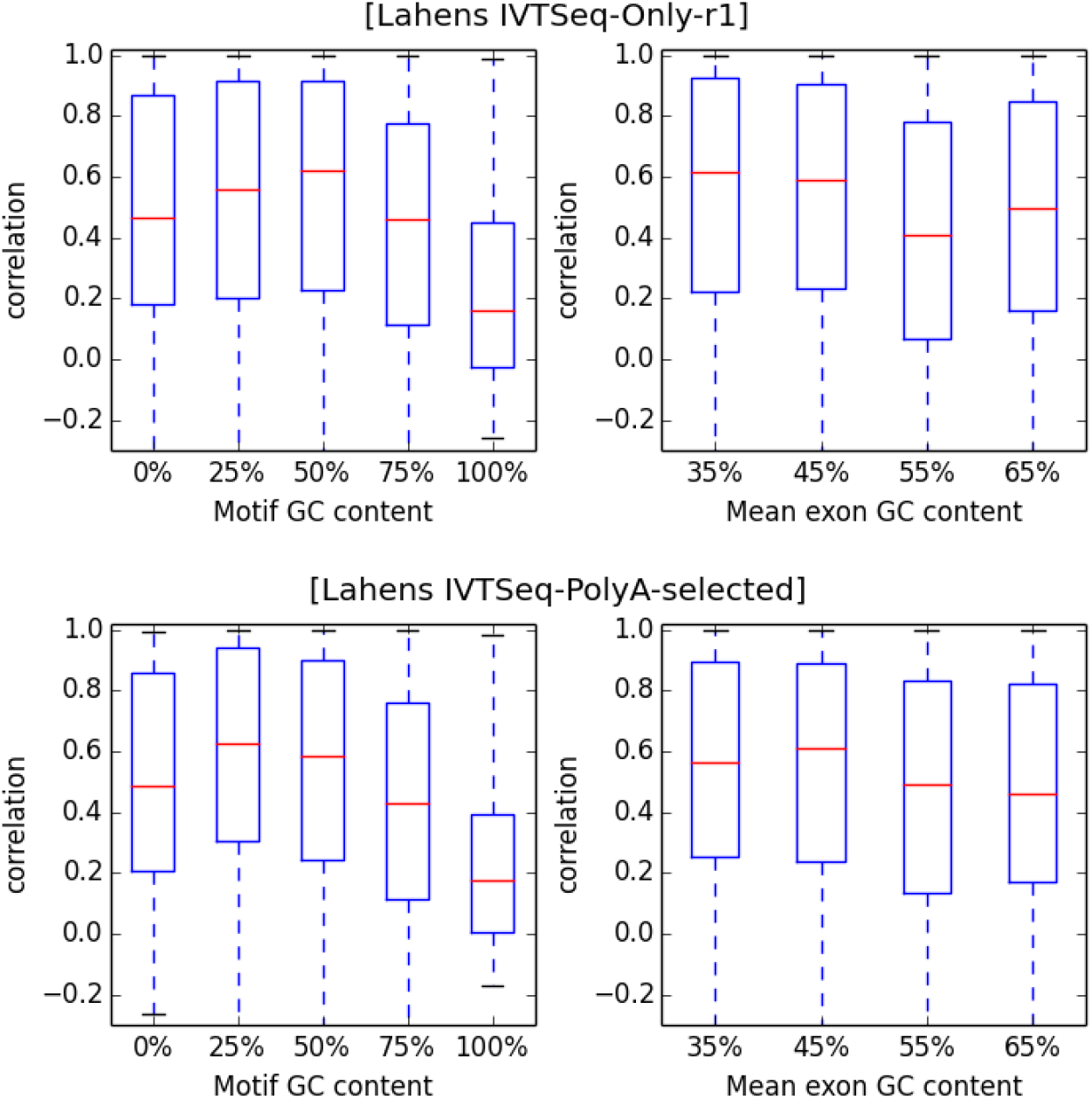
Correlation (Pearson’s) as a function of *4-mer* motif and exon GC content for *H. sapiens* for the two IVT samples that had different RNA selection protocols applied: Top) *IVT-Only* sample, which underwent ribosomal depletion, and B) *IVT-PolyA* which underwent PolyA selection.

### 3.2 Analysis of Wild-type and Mutant-r2 *D. melanogaster*

Using our method, we have also analysed the whole transcriptomes of two Drosophila Fruitfly (species *D. melanogaster*) which only differ by mutation *gl[60j]* in the eye-antennal disc [39]. Overall correlations for both wild-type and mutant-r2 are given in Table 4. We observe much lower correlations for the mutant-r2 than in wild-type — the difference is especially marked when comparing the motifs with the highest 10 correlation in both samples.

**Table 4:**
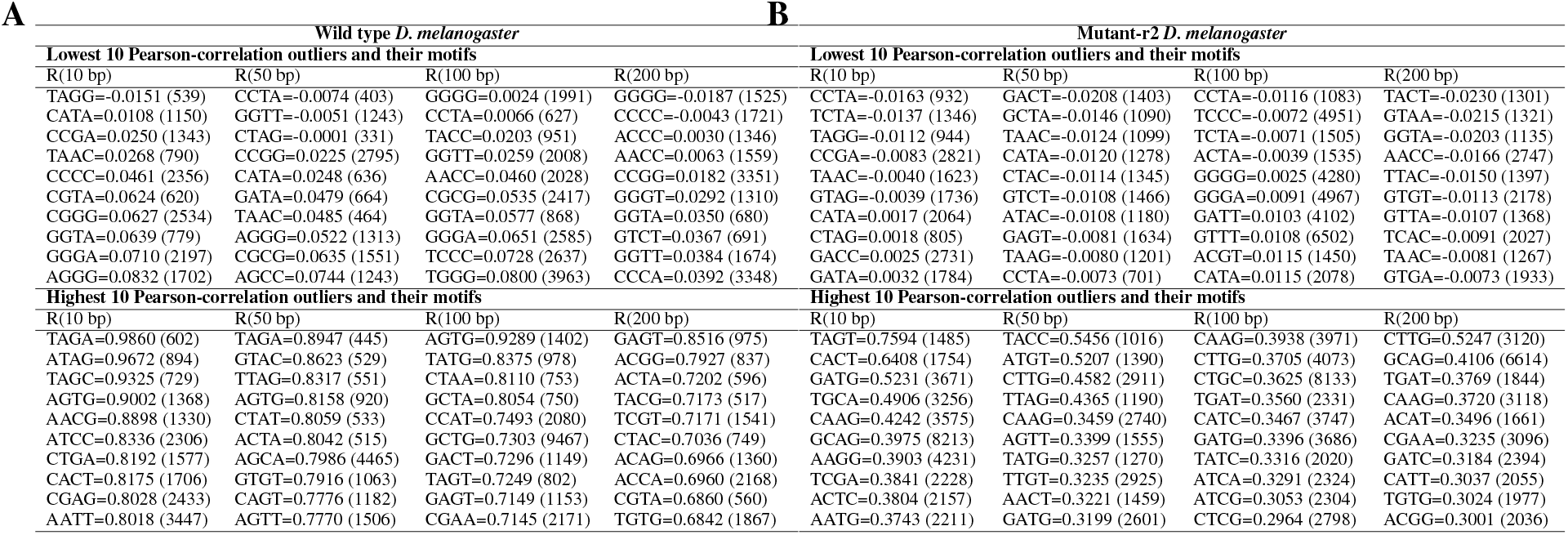
*D. melanogaster* Pearson correlation co-efficient outliers (top ten and lowest ten) for different *intra-exon 4-mer* motif sequence pairs at 10, 50, 100 and 200 bp spacings. A) Wild-type B) Mutant-r2. Sample sizes are given in parenthesis. Replicates are given in Supplementary Table S2.

#### 3.2.1 GC content of the replicates

In order to examine the effect of GC content on the distribution of mapped reads, we have plotted intra-exon *4-mer* motif-pair correlations as a function of both motif and exon GC content, for both *D. melanogaster* datasets (Figure 3). In the wild-type dataset we observe notable variation in the correlation as a function of GC content of the motif, whereas no variation in the correlation is observed as a function of the mean exon GC content. This indicates that the GC content of the motif has an effect (causing a deviation in the distribution of mapped reads to an exon) rather than the overall GC content. In the mutant-r2 dataset, no variation is observed.

**Figure 3:**
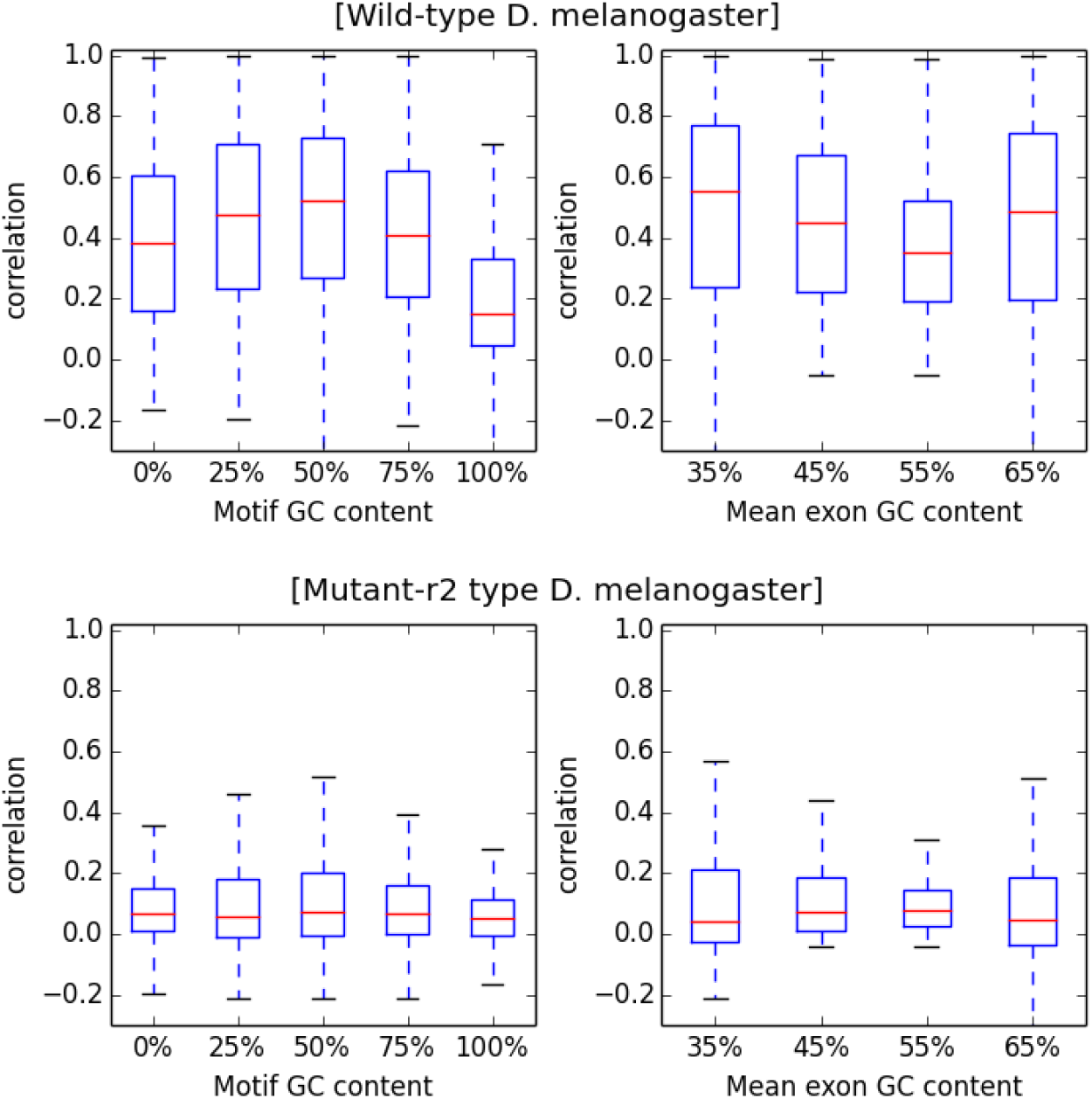
Correlation (Pearson’s) as a function of *4-mer* motif and exon GC content in both wild and mutant-r2 *D. melanogaster* transcriptomes. NB: The replicates are given in Supplementary Figure S4

**Figure 4:**
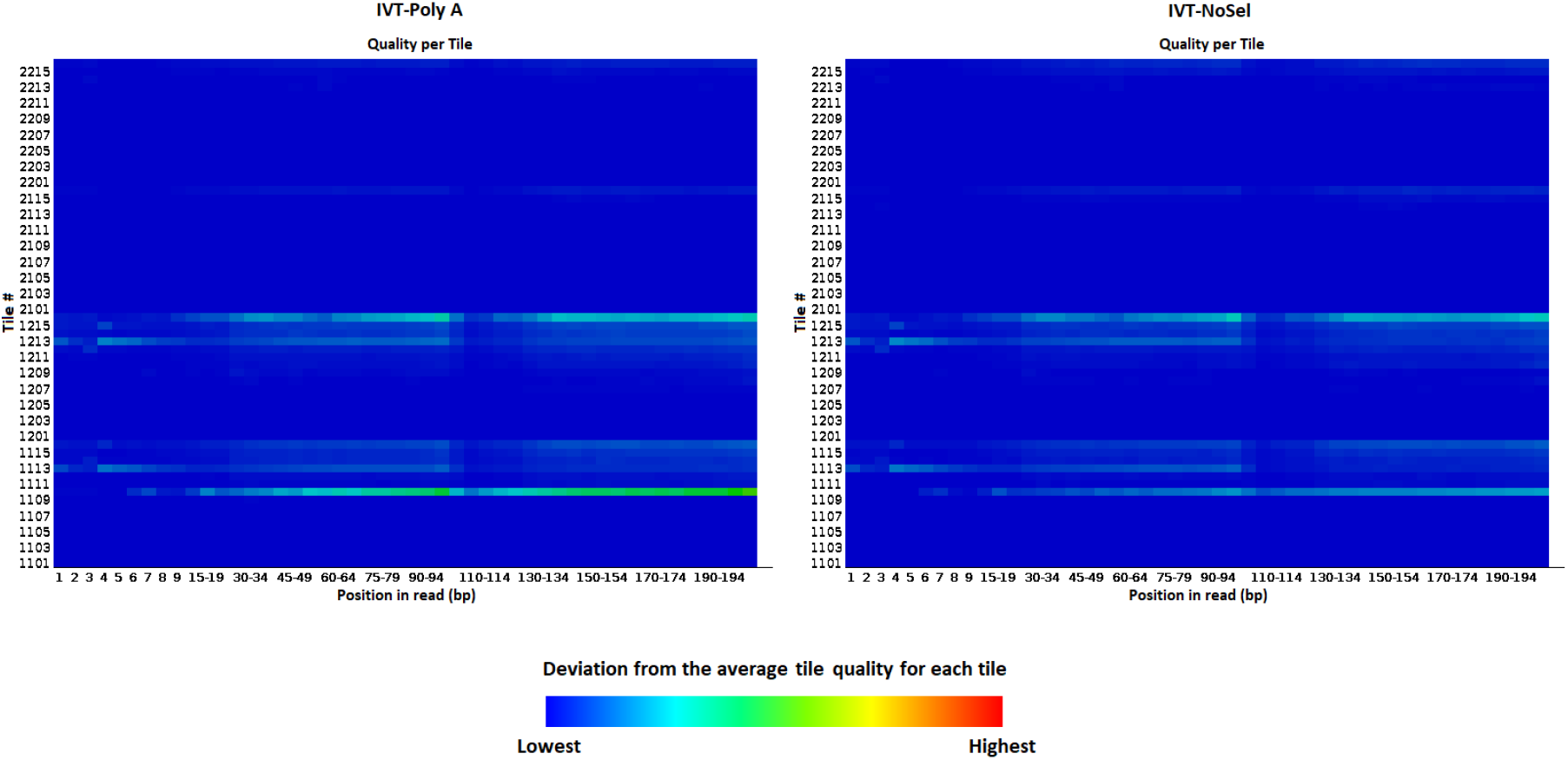
Heatmaps of the Sequencing *per-tile* Means, obtained from FastQC analysis for the two RNA-selection *H. sapiens* IVT RNA-Seq samples analysed — sequencing tile information was not present in the *IVT-Plasmids* sample. Left) *IVT-PolyA* Right) *IVT-NoSel*.

### 3.3 FastQC and BamQC — Quality Control analysis of *D. melanogaster* dataset

Given there is no biological precedent for the differences observed between the wild and mutant species, and that the low correlations are seen across the mutant replicates, independent of both spacing and GC content, the source of variance is likely to be technical in origin. With this in mind, in order to see if the source of variance between the *D. melanogaster* datasets could be identified by means of Quality Control (QC) checks, we ran Qualimap [43] on all of the *D. melanogaster* samples and replicates, and then ran FastQC [30] on all of the datasets we analysed. In this way, we worked our way backwards through the analysis (i.e. working upstream from our analysis method), starting with the *D. melanogaster* datasets. Firstly, in order to see if there were any alignment or coverage issues in the mutant dataset, we ran Qualimap to analyse the source BAM (Binary Alignment files) that the SAM files were directly derived from. Next, in order to analyse the raw unaligned reads files (Fasta files) for any sequencing issues, we ran FastQC on all of the RNA-Seq replicates and samples, for all datasets that we have analysed in this research.

When we compare the BamQC analysis for the wild and mutant types (a summary of the results is given in Table 5, there is not much difference between them in terms of the mapping quality, nucleotide frequencies and GC content. However, there is an increase in variance in the coverage of the mutant *D. melanogaster* dataset as indicated by the slightly higher standard deviation (93.045 vs. 80.229). To look into this further, we looked into how coverage across chromosomes in the two samples was reported in the BamQC report, this is plotted in Figure 7. This shows substantial variation in coverage between different chromosomes that reads were mapped to in the organism. However a similar pattern is observed in all of the samples and thus cannot account for the differences in *4-mer* correlations that we observe.

**Table 5:** BamQC analysis summary for (A) wild-type and (B) mutant-r2 Drosophila.

| A | Wild-type *D. melanogaster* |  |
| --- | --- | --- |
|  | Parameter | Result |
|  | number of reads | 12,960,778 |
|  | number of mapped reads | 12,960,778 (100%) |
|  | number of mapped bases | 554,135,221 bp |
|  | number of sequenced bases | 442,559,999 bp |
|  | mean mapping quality | 135.086 |
|  | mean coverage data | 3.284X |
|  | std coverage data | 80.2288X |
| B | Mutant-r2 *D. melanogaster* |  |
|  | Parameter | Result |
|  | number of reads | 15,099,081 |
|  | number of mapped reads | 15,099,081 (100%) |
|  | number of mapped bases | 656,286,624 bp |
|  | number of sequenced bases | 530,325,061 bp |
|  | mean mapping quality | 139.9086 |
|  | mean coverage data | 3.8894X |
|  | std coverage data | 93.0446X |

| A | Base composition / GC content |  |
| --- | --- | --- |
|  | Parameter | Result |
|  | number of A's | 114,904,069 bp (25.96%) |
|  | number of C's | 107,246,603 bp (24.23%) |
|  | number of T's | 115,649,248 bp (26.13%) |
|  | number of G's | 104,760,079 bp (23.67%) |
|  | number of N's | 111,575,222 bp (25.21%) |
|  | GC percentage | 47.9% |

| B | Base composition / GC content |  |
| --- | --- | --- |
|  | Parameter | Result |
|  | number of A's | 136,501,323 bp (25.74%) |
|  | number of C's | 129,898,946 bp (24.49%) |
|  | number of T's | 137,081,079 bp (25.85%) |
|  | number of G's | 126,843,713 bp (23.92%) |
|  | number of N's | 125,961,563 bp (23.75%) |
|  | GC percentage | 48.41% |

In comparing the FastQC analysis results for the datasets, we found that the *per tile sequencing quality* heatmaps were radically different for the mutant and wild type datasets (Figure 5). These depict deviations from the average tile quality within areas of flowcells on the sequencing apparatus (the explicit values underlying the heatmaps are referred to as the per-tile Means) [44]. In the wild-type replicates, these heatmaps were plain and uniform, whereas both of the mutant replicates showed considerable variation from the Means. There were no differences in any of the other FastQC analysis parameters measured between the wild and mutant datasets. In order to investigate this effect further, and to contrast it with the IVT dataset samples, we plotted the distribution of per-tile Means as both a histogram and a density plot (Figure 6) for all the samples in all of the datasets we have analysed. In Figure 6, we observe that the distributions in the IVT and wild type datasets show the least deviation from the Mean tile qualities. The mutant *D. melanogaster* datasets show considerable deviation from the Mean tile qualities — this accounts for the low correlations seen across the mutant replicates. By computing *intra-exon* motif-pair correlations we have observed that *sequencing errors occurring in flowcell tiles result in widespread deviations from the uniformity of mapped reads across exons, and are independent of sequence or GC content*.

**Figure 5:**
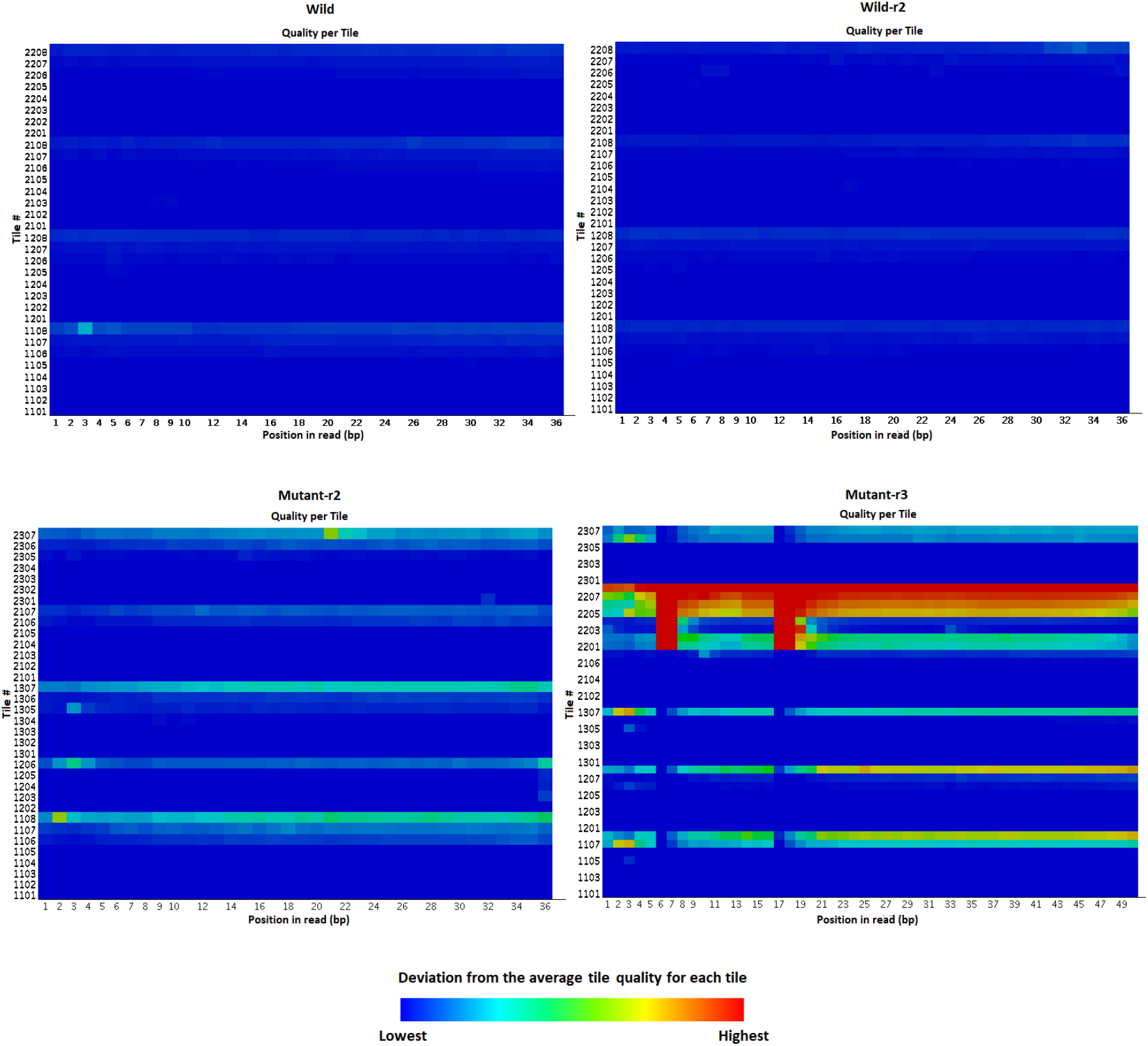
Heatmaps of the Sequencing *per-tile* Means, obtained from FastQC analysis for all *D. melanogaster* RNA-Seq samples and their replicates analysed Top) Wild-type replicates Bottom) Mutant replicates.

**Figure 6:**
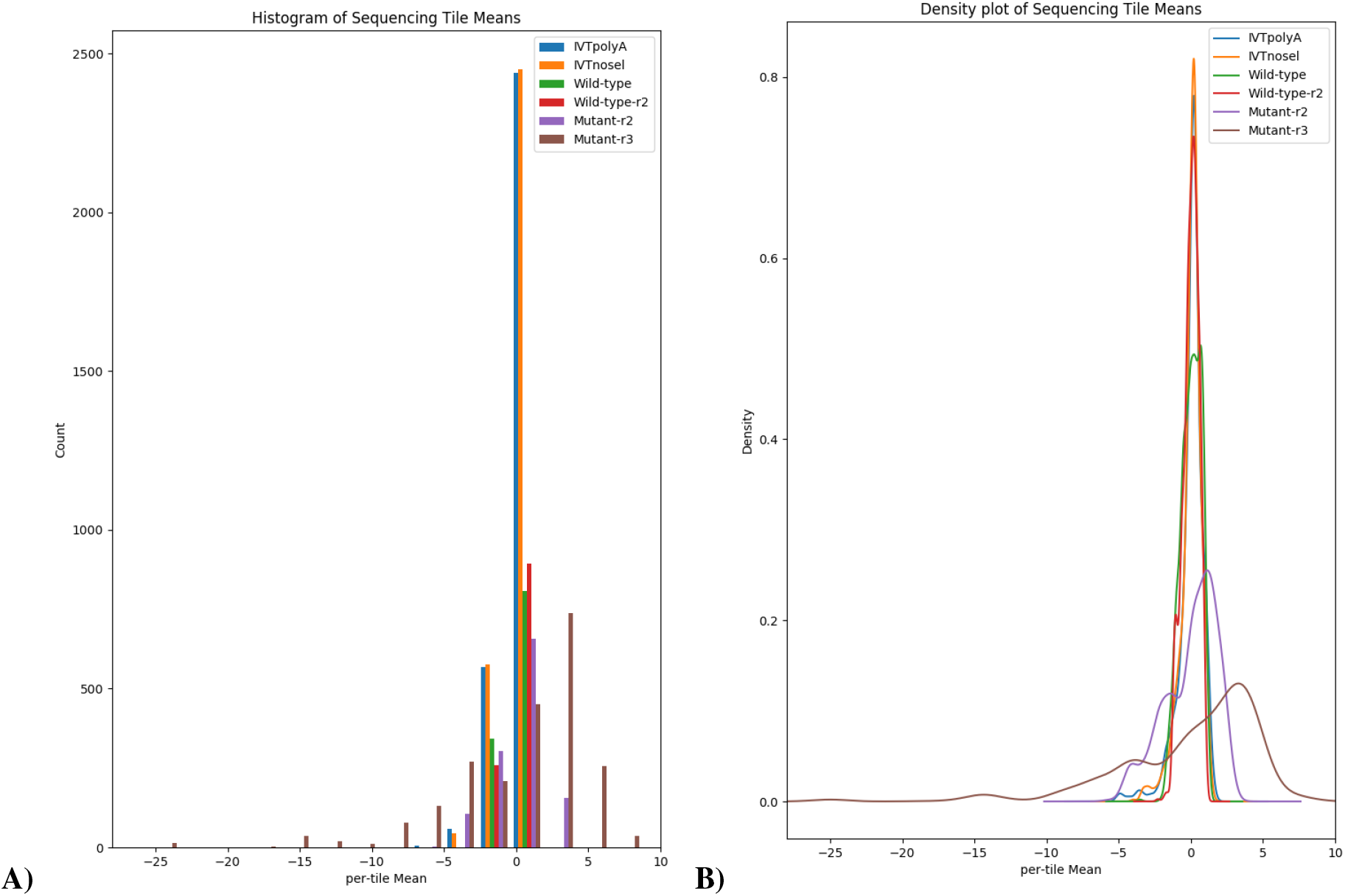
Visualisation of the distributions of the Sequencing *per-tile* Means, obtained from FastQC analysis for all RNA-Seq samples analysed (species *D. melanogaster* and *H. sapiens*). A) Histogram B) Density plot. The distributions in the IVT datasets, and both wild type replicates, show that less of the tiles in the flowcells of the sequencing apparatus deviate from the *per-tile* Means. This is indicated by the tallest, narrowest peaks, centered around 0, for the IVTPolyA dataset (Blue) followed by IVTnosel (Orange), and then the first wild-type replicate (Red). The second wild-type replicate (Green) was the third tallest peak, and showed a reasonably good narrow distribution of the data about the *per-tile* Means, albeit lower than the IVT datasets and first wild type replicate. The two mutant replicates, however, show very wide distributions about the *per-tile* Means in the flowcells, as depicted by the shorter, wider curves (Purple and Brown) respectively. NB: The *IVT-Plasmids* sample did not carry sequencing tile information.

**Figure 7:**
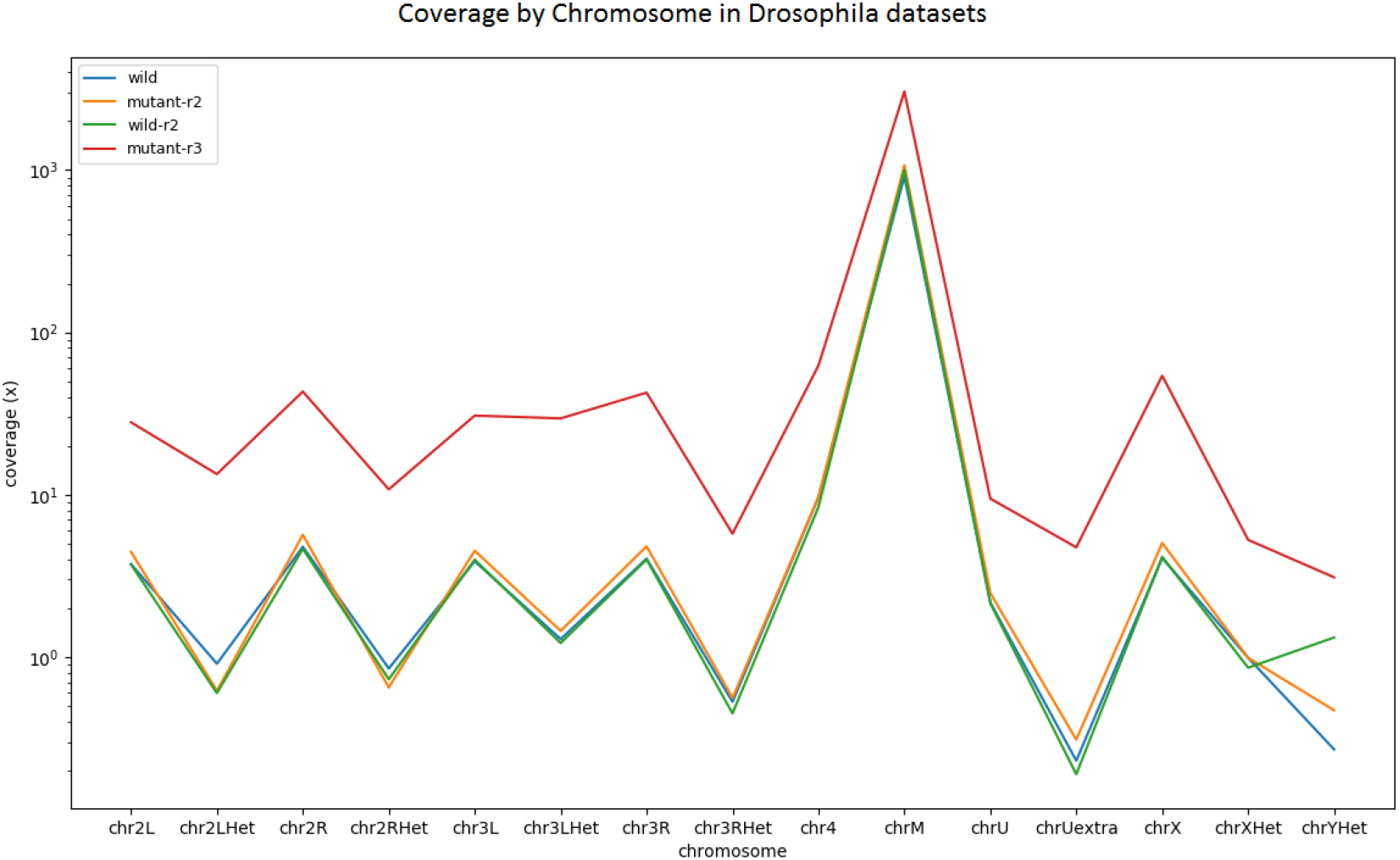
Plot of chromosome coverage, obtained from BamQC analysis results for wild-type and mutant-r2 Drosophila.

## 4 Discussion

We have analysed RNA-Seq samples from two species: *H. sapiens* (MGC plasmids) and *D. melanogaster* (transcriptomes). The IVT dataset was produced in a highly controlled manner — we have analysed three *H.sapiens* samples from this, two were subjected to mRNA selection prior to RNA-Seq, and the other was not. The IVT samples contain fewer reads than the *D. melanogaster* samples in which the wild and mutant-r2 types, comprise approximately 12.9 M and 15.0 M reads respectively, whilst the *IVT-Only* sample has 0.406 M reads, the *IVT-PolyA* sample has 0.397 M reads and the *IVT-Plasmids* 0.181 M reads. Although these are comparatively small numbers of reads, the IVT samples have the *intra-exon* bias within them, as indicated by coverage of reads mapping to the source MGC plasmids used, well quantified. From the box and whisker plots of the two IVT samples that underwent mRNA selection RNA-Seq protocols (Ribosomal depletion and Poly-A selection), we observe a dependence of *intra-exon* motif pair correlation on motif GC content. This effect is also seen in the wild-type replicates of the *D. melanogaster* samples, but not in the *IVT-Plasmids* sample which is not subjected to mRNA selection. The mutant D. melanogaster data has very low overall correlations and hence this pattern is not seen here.

### 4.1 Dependence of *intra-exon* correlation on GC content appears to be due to mRNA selection

The *intra-exon* motif correlations as a function of both GC content parameters are much higher in the *IVT-Plasmids* mRNA *selection free* RNA-Seq sample than in the other RNA-Seq samples that we analysed that did undergo mRNA selection: both ribosomal depletion (*IVT-Only*) and PolyA selection (*IVT-polyA* and wild-type). Furthermore, both of the *H. sapiens* and wild-type *D. melanogaster* samples that underwent mRNA selection in the RNA-Seq process had slightly lower correlations than the H. sapiens *IVT-Plasmids* sample which did not, suggesting this is likely of technical origin. Importantly, all the samples from all of the datasets we analysed were sequenced on the same platform — Illumina HiSeq 2000 (as detailed in Table 1).

As the dependence on overall GC concentration is not observed in the *IVT-Plasmids* control sample, but is observed in RNA-Seq samples that underwent mRNA selection, we can exclude platform-specific sequencing bias as a source of this effect in the IVT samples. This suggests that not only do mRNA selection protocols result in bias in the distribution of mapped RNA-Seq reads as Lahens et al. demonstrated [24], but that *mRNA selection is also responsible for the dependence of correlation on GC content* — we have observed this in all of the RNA-Seq samples that underwent mRNA selection across both *H. sapiens* and *D. melanogaster* species. Risso et al. also noted motif-specific GC effects, manifesting as deviations from uniform read distribution, which they attribute these problematic motifs being underrepresented [45]. However, the GC effect we observe occurs in both the wild type replicates, which have large sample sizes (A of Table 4), as well as in the *IVT-polyA* and *IVTnosel* mRNA selection datasets, which have low sample sizes (due to the controlled way in which the samples were prepared) (Table 3). Furthermore, the numbers of counts for high GC content motifs, as indicated by the highest outliers (Table 4), is of the same order as other motifs. The dependence of *intra-exon* correlations on the GC content of the motifs appears to be due to mRNA selection methods, which are routinely employed in RNA-Seq experiments, and are known to introduce bias [46, 47, 24].

## 5 Conclusion

In this work, we have applied our novel *k-mer* based analysis method that allows deep investigation into RNA-Seq read data at the exon level, and quantifies sequence-specific deviations in the uniformity of the distribution of mapped reads to a reference genome. The method we have applied uses distributed computing to count reads overlapping *4-mer* motifs exhaustively (AAAA through to GGGG) at different regions of the same exon. The assumption is that, when there is no deviation in the distribution of mapped reads, these counts should be correlated. We have demonstrated that the correlations we have computed correspond to mean tile data in the samples from the two datasets we analysed, and propose that this be used when mean tile data is not available and therefore requires only the raw short-read data alone. We have also observed that extremely poor correlations are an indication of technical sources of bias, such as sequencing flowcell tile errors or batch-effects. This is important work because i) gene expression studies rely on abundance estimates of RNA transcripts that can be hampered by deviations in the uniformity of read distribution ii) the lack of annotation in experiments deposited at public repositories, such as the SRA and GEO, compound the challenges in characterising bias in RNA-Seq data, and iii) our method is scalable and can, therefore, be applied to analyse the large datasets present in these vast data repositories. As of August 2020 (the time of writing) the SRA alone contains more than 43 peta bases (43.390×10^15^) of sequencing data, that is in excess of 5 terabytes [48]. In light of this increase in throughput, the relatively young age of RNA-Seq (hence not all bias has been characterised), and the trend towards data-driven science, more work is needed to develop QC measures to ensure the integrity of this data.

## 6 Acknowledgements

The authors wish to thank Eszter Ábrahám for proofreading the manuscript.

## Notes

https://github.com/jamie-alnasir/hercules

### Competing Interest Statement

The authors have declared no competing interest.

## References

[1] Steven Soper. Dna sequencing using fluorescence detection. volume 230, pages 1350–1354. 1985.

[2] Lloyd M Smith, Jane Z Sanders, Robert J Kaiser, Peter Hughes, Chris Dodd, Charles R Connell, Cheryl Heiner, SB Kent, and Leroy E Hood. Fluorescence detection in automated dna sequence analysis. Nature, 321(6071):674–679, 1985.

[3] James M Prober, George L Trainor, Rudy J Dam, Frank W Hobbs, Charles W Robertson, Robert J Zagursky, Anthony J Cocuzza, Mark A Jensen, and Kirk Baumeister. A system for rapid dna sequencing with fluorescent chain-terminating dideoxynucleotides. Science, 238(4825):336–341, 1987.

[4] Elaine R Mardis. Anticipating the 1,000 dollar genome. Genome biology, 7(7):112, jan 2006.

[5] Jay Shendure and Hanlee Ji. Next-generation dna sequencing. Nature biotechnology, 26(10):1135–1145, 2008.

[6] Michael L Metzker. Sequencing technologies - the next generation. Nature reviews. Genetics, 11(1):31–46, jan 2010.

[7] Nayanah Siva. Uk gears up to decode 100 000 genomes from nhs patients. The Lancet, 385(9963):103–104, 2015.

[8] James Gallagher, BBC. DNA project ’to make UK world genetic research leader’. http://www.bbc.co.uk/news/health-28488313, 2014. [Online; accessed 21-January-2019].

[9] Clarke, T. and Begley S., Reuters. 1000 Genomes Project Releases Data from Pilot Projects on Path to Providing Database for 2,500 Human Genomes - Freely available data supporting next generation of human genetic research. http://www.reuters.com/article/us-usa-obama-precisionmedicine-idUSKBN0L313R20150130, 2015. [Online; accessed 02-February-2019].

[10] David Cyranoski. China’s bid to be a dna superpower. Nature, 534(7608):462–463, 2016.

[11] NIH. Scientists Form International Cancer Genome Consortium. https://www.nih.gov/news-events/news-releases/scientists-form-international-cancer-genome-consortium, 2008. [Online; accessed 24-November-2019].

[12] Steve Russell, Lisa A Meadows, and Roslin R Russell. Microarray technology in practice. Academic Press, 2008.

[13] Zhong Wang, Mark Gerstein, and Michael Snyder. Rna-seq: a revolutionary tool for transcriptomics. Nature reviews genetics, 10(1):57–63, 2009.

[14] Kimberly R Kukurba and Stephen B Montgomery. Rna sequencing and analysis. Cold Spring Harbor Protocols, 2015(11):pdb–top084970, 2015.

[15] Juw Won Park, Collin Tokheim, Shihao Shen, and Yi Xing. Identifying differential alternative splicing events from rna sequencing data using rnaseq-mats. Deep Sequencing Data Analysis, pages 171–179, 2013.

[16] Ruolin Liu, Ann E Loraine, and Julie A Dickerson. Comparisons of computational methods for differential alternative splicing detection using rna-seq in plant systems. BMC bioinformatics, 15(1):1, 2014.

[17] Ali Mortazavi, Brian A Williams, Kenneth McCue, Lorian Schaeffer, and Barbara Wold. Mapping and quantifying mammalian transcriptomes by rna-seq. Nature methods, 5(7):621–628, 2008.

[18] Manuel Garber, Manfred G Grabherr, Mitchell Guttman, and Cole Trapnell. Computational methods for transcriptome annotation and quantification using rna-seq. Nature methods, 8(6):469–477, 2011.

[19] Tal Raz, Philipp Kapranov, Doron Lipson, Stan Letovsky, Patrice M Milos, and John F Thompson. Protocol dependence of sequencing-based gene expression measurements. PloS one, 6(5):e19287, 2011.

[20] Fatih Ozsolak, Adam R Platt, Dan R Jones, Jeffrey G Reifenberger, Lauryn E Sass, Peter McInerney, John F Thompson, Jayson Bowers, Mirna Jarosz, and Patrice M Milos. Direct rna sequencing. Nature, 461(7265):814–818, 2009.

[21] Frazer Meacham, Dario Boffelli, Joseph Dhahbi, David I K Martin, Meromit Singer, and Lior Pachter. Identification and correction of systematic error in high-throughput sequence data. BMC bioinformatics, 12(1):451, jan 2011.

[22] Juliane C Dohm, Claudio Lottaz, Tatiana Borodina, and Heinz Himmelbauer. Substantial biases in ultra-short read data sets from high-throughput DNA sequencing. Nucleic acids research, 36(16):e105, sep 2008.

[23] Jamie Alnasir and Hugh P Shanahan. Investigation into the annotation of protocol sequencing steps in the sequence read archive. GigaScience, 4:23, 2015.

[24] Nicholas F Lahens, Ibrahim Halil Kavakli, Ray Zhang, Katharina Hayer, Michael B Black, Hannah Dueck, Angel Pizarro, Junhyong Kim, Rafael Irizarry, Russell S Thomas, et al. Ivt-seq reveals extreme bias in rna sequencing. Genome biology, 15(6):R86, 2014.

[25] Adam Roberts, Cole Trapnell, Julie Donaghey, John L Rinn, and Lior Pachter. Improving RNA-Seq expression estimates by correcting for fragment bias. Genome biology, 12(3):R22, 2011.

[26] Kasper D Hansen, Steven E Brenner, and Sandrine Dudoit. Biases in Illumina transcriptome sequencing caused by random hexamer priming. Nucleic acids research, 38(12):e131, jul 2010.

[27] Rasko Leinonen, Hideaki Sugawara, and Martin Shumway. The sequence read archive. Nucleic acids research, 39(Database issue):D19–21, jan 2011.

[28] R. Edgar. Gene Expression Omnibus: NCBI gene expression and hybridization array data repository. Nucleic Acids Research, 30(1):207–210, jan 2002.

[29] Takeru Nakazato and Ohta et al. Experimental Design-Based Functional Mining and Characterization of High-Throughput Sequencing Data in the Sequence Read Archive. PLoS ONE, 8(10):e77910, oct 2013.

[30] Simon Andrews et al. Fastqc: a quality control tool for high throughput sequence data, 2010.

[31] Jamie Alnasir and Hugh P Shanahan. A novel method to detect bias in short read ngs data. Journal of integrative bioinformatics, 14(3), 2017.

[32] Zachary D. Stephens, Skylar Y. Lee, Faraz Faghri, Roy H. Campbell, Chengxiang Zhai, Miles J. Efron, Ravishankar Iyer, Michael C. Schatz, Saurabh Sinha, and Gene E. Robinson. Big data: Astronomical or genomical? PLoS Biol, 13(7):1–11, 07 2015.

[33] J Alnasir. Source code and results data for Transcriptomics Analysis System (Hercules). https://github.com/jamie-alnasir/hercules, 2018. [Online; accessed 20-August-2020].

[34] Gary Temple, Daniela S Gerhard, Rebekah Rasooly, Elise A Feingold, Peter J Good, Cristen Robinson, Allison Mandich, Jeffrey G Derge, Jeanne Lewis, Debonny Shoaf, et al. The completion of the mammalian gene collection (mgc). Genome research, 19(12):2324–2333, 2009.

[35] Yen-Chun Chen, Tsunglin Liu, Chun-Hui Yu, Tzen-Yuh Chiang, and Chi-Chuan Hwang. Effects of gc bias in next-generation-sequencing data on de novo genome assembly. PloS one, 8(4):e62856, 2013.

[36] Farhat Naureen Memon, Anne M Owen, Olivia Sanchez-Graillet, Graham JG Upton, and Andrew P Harrison. Identifying the impact of g-quadruplexes on affymetrix 3’arrays using cloud computing. Journal of integrative bioinformatics, 7(111), 2010.

[37] Zhang R Hayer K Black MB Dueck H Pizarro A Kim J Irizarry R Thomas RS Grant GR Hogenesch JB Lahens NF, Kavakli IH. GSE50445: IVT-seq reveals extreme bias in RNA-sequencing. https://www.ncbi.nlm.nih.gov/geo/query/acc.cgi?acc=GSE50445, 2012. [Online; accessed 01-November-2019].

[38] Stein Aerts. GEO-GSE39781: RNA-seq in wild-type and glass mutant eye-antennal discs in Drosophila melanogaster. https://www.ncbi.nlm.nih.gov/geo/query/acc.cgi?acc=GSE39781, 2012. [Online; accessed 27-March-2019].

[39] Marina Naval-Sánchez, Delphine Potier, Lotte Haagen, Máximo Sánchez, Sebastian Munck, Bram Van de Sande, Fernando Casares, Valerie Christiaens, and Stein Aerts. Comparative motif discovery combined with comparative transcriptomics yields accurate targetome and enhancer predictions. Genome research, 23(1):74–88, 2013.

[40] A Gordon and GJ Hannon. Fastx-toolkit. FASTQ/A short-reads preprocessing tools (unpublished) http://hannonlab.cshl.edu/fastx_toolkit, 2010.

[41] Cole Trapnell, Lior Pachter, and Steven L Salzberg. Tophat: discovering splice junctions with rna-seq. Bioinformatics, 25(9):1105–1111, 2009.

[42] L Sian Gramates, Steven J Marygold, Gilberto dos Santos, Jose-Maria Urbano, Giulia Antonazzo, Beverley B Matthews, Alix J Rey, Christopher J Tabone, Madeline A Crosby, David B Emmert, et al. Flybase at 25: looking to the future. Nucleic Acids Research, page gkw1016, 2016.

[43] Fernando García-Alcalde, Konstantin Okonechnikov, José Carbonell, Luis M Cruz, Stefan Götz, Sonia Tarazona, Joaquín Dopazo, Thomas F Meyer, and Ana Conesa. Qualimap: evaluating next-generation sequencing alignment data. Bioinformatics, 28(20):2678–2679, 2012.

[44] Babraham Institute. FastQC documentation - per-tile sequencing quality. https://www.bioinformatics.babraham.ac.uk/projects/fastqc/Help/3AnalysisModules/12PerTileSequenceQuality.html, 2015. [Online; accessed 03-Dec-2019].

[45] Davide Risso, Katja Schwartz, Gavin Sherlock, and Sandrine Dudoit. Gc-content normalization for rna-seq data. BMC bioinformatics, 12(1):480, 2011.

[46] Douglas Kyung Nam, Sanggyu Lee, Guolin Zhou, Xiaohong Cao, Clarence Wang, Terry Clark, Jianjun Chen, Janet D Rowley, and San Ming Wang. Oligo (dt) primer generates a high frequency of truncated cdnas through internal poly (a) priming during reverse transcription. Proceedings of the National Academy of Sciences, 99(9):6152–6156, 2002.

[47] Zhao Zhang, William E Theurkauf, Zhiping Weng, and Phillip D Zamore. Strand-specific libraries for high throughput rna sequencing (rna-seq) prepared without poly (a) selection. Silence, 3(1):9, 2012.

[48] Sequence Read Archive. Overview of the Sequence Read Archive (SRA). https://trace.ncbi.nlm.nih.gov/Traces/sra/sra.cgi/, 2017. [Online; accessed 3-January-2019].

