## Supplementary Information for "Intra-exon motif correlations as a proxy measure for mean per-tile sequence quality data in RNA-Seq"

### 1 Testing - Synthetic transcriptomics read data

To validate our method for quantifying sequence-specific deviations in the read distribution of mapped reads we created a synthetic transcriptome. This consists of an artificial GTF genome annotation of artificial chromosomes and their features together with a SAM reads file of artificial reads. The SAM reads file was constructed by randomly generating a 50,000 bp sequence template seed in the form of a string of 50,000 characters. The base composition of this seed is shown in Table S1 below:

| **Synthetic seed** | |
|---|---|
| **Base** | **Count** |
| Adenine (A) | 12,558 |
| Thymine (T) | 12,662 |
| Guanine (G) | 12,297 |
| Cytosine (C) | 12,483 |
| **Total bases** | 50000 |
| **GC content** | 49.6% |

Table S1: Nucleotide composition of synthetic sequence seed fragment used to generate synthetic transcriptomic reads for calibration of the algorithm.

Artificial "synthetic" reads were then created by generating random start and random end positions from which reads were constructed by copying the sequence from the seed. We deemed it unnecessary to derive complementary reads from the template as the reference genome is not necessary, and therefore we chose to generate corresponding CIGAR strings for each read which were contiguous alignment matches of the same length as the artificial read sequences. Each sequence read has the same length of its exon feature, but has varying number of copies - this represents uniform distribution of reads but at different levels of expression.

In applying our analysis method to this synthetic dataset we expect to observe only perfect Pearson correlations for any *intra-exon* sequence motif-pairs found within the inspected reads.

In real transcriptomics data the distribution of reads means that motif-pairs spaced farther apart will show poorer correlation. In the synthetic reads this should not be the case because the reads are generated as contiguous reads which are of the same length as the exon. In order to verify this we have applied our analysis method to this synthetic dataset and plotted the results as a scattermatrix plot of correlation $\rho$ ($r^2$), partitioned by spacing $d$, motif GC $gm$ and mean GC content of reads in the exon $\overline{ge}$ (Figure S1). We can see in the scattermatrix plot that all of the *intra-exon* motif pairs are correlated at +1.0 (postive, perfect correlation) for all parameters. It also shows that there is no dependence of correlation on GC content. This is as we would expect from a synthetic randomly generated transcriptome in which there have been no molecular or sequencing processes applied.

### 2 Supplementary Results

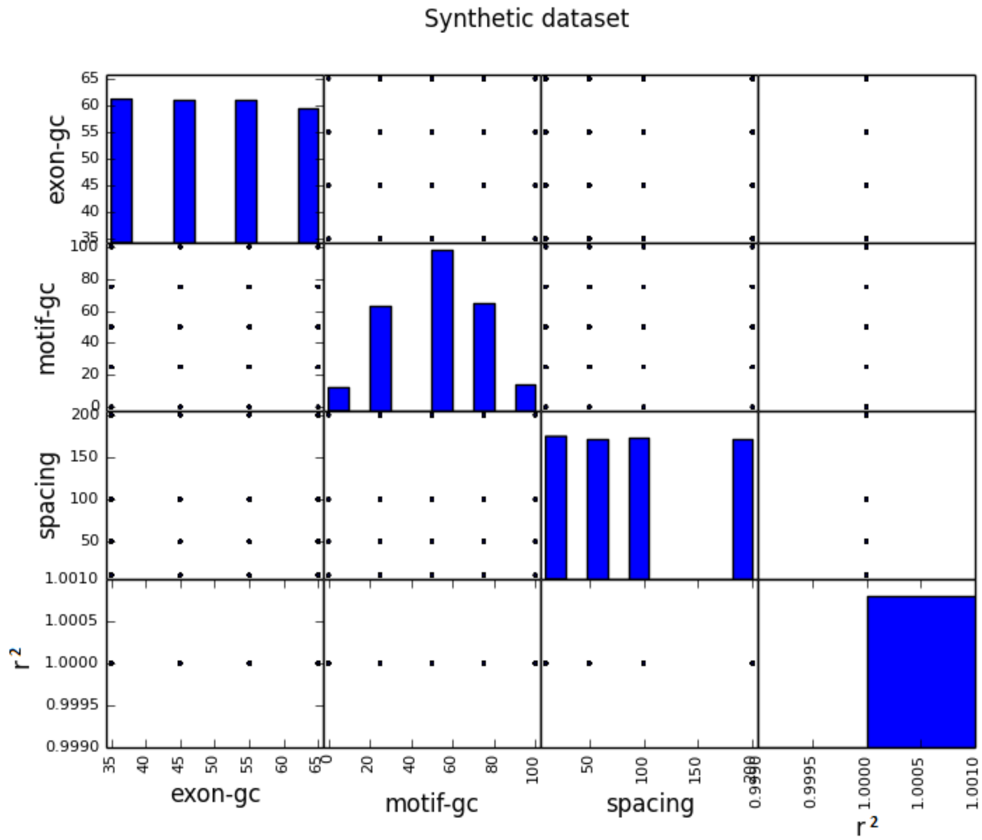


Figure S1: Comparison of correlations as a function of Motif and Exon GC content in the synthetic transcriptome. The distribution of correlations (lower right-most sub-plot) shows that the *intra-exon k-mer* pairs are perfectly correlated (score of +1). Distributions are shown horizontally by bar charts (in blue).

**A**

| **Wild-r2 type *D. melanogaster*** | | | |
|---|---|---|---|
| **Lowest 10 Pearson-correlation outliers and their motifs** | | | |
| R(10 bp) | R(50 bp) | R(100 bp) | R(200 bp) |
| TAGG=-0.0154 (534) | CCTA=-0.0078 (392) | GGGG=0.0023 (1977) | GGGG=-0.0188 (1507) |
| CATA=0.0100 (1141) | CTAG=0.0005 (320) | CCTA=0.0065 (611) | CCCC=-0.0045 (1699) |
| CCGA=0.0248 (1337) | GGTT=0.0173 (757) | CGCG=0.0533 (2398) | ACCC=0.0030 (1332) |
| TAAC=0.0267 (781) | CATA=0.0204 (630) | GGGA=0.0648 (2538) | CCGG=0.0185 (3330) |
| CCCC=0.0460 (2346) | CCGG=0.0224 (2776) | AACC=0.0713 (1206) | GGGT=0.0288 (1286) |
| CGTA=0.0622 (613) | GATA=0.0479 (662) | TCCC=0.0736 (2599) | GTCT=0.0370 (684) |
| CGGG=0.0625 (2519) | TAAC=0.0484 (456) | TGGG=0.0797 (3909) | CCCA=0.0391 (3305) |
| GGGA=0.0713 (2185) | AGGG=0.0520 (1302) | CGGG=0.0904 (3318) | AACC=0.0425 (957) |
| AGGG=0.0831 (1690) | CGCG=0.0629 (1524) | TCTC=0.0930 (1669) | AGGG=0.0479 (1509) |
| CGTT=0.0844 (1242) | AGCC=0.0747 (1237) | CATA=0.1012 (1018) | ATAG=0.0550 (532) |
| **Highest 10 Pearson-correlation outliers and their motifs** | | | |
| R(10 bp) | R(50 bp) | R(100 bp) | R(200 bp) |
| TAGA=0.9860 (600) | TAGA=0.9033 (440) | TATG=0.8378 (964) | GAGT=0.8517 (961) |
| ATAG=0.9672 (892) | TACC=0.8850 (336) | GCTA=0.8293 (743) | ACGG=0.7927 (836) |
| TAGC=0.9325 (727) | GTAC=0.8623 (523) | CTAA=0.8113 (738) | ACTA=0.7205 (588) |
| AACG=0.8898 (1322) | GGTA=0.8370 (357) | TAGT=0.7988 (787) | TCGT=0.7189 (1534) |
| AGTG=0.8618 (1575) | TTAG=0.8317 (544) | CCAT=0.7495 (2063) | TACG=0.7173 (514) |
| ATCC=0.8336 (2296) | AGTG=0.8219 (1072) | GCTG=0.7304 (9423) | CTAC=0.7047 (740) |
| CTGA=0.8193 (1555) | CTAT=0.8069 (518) | AGTG=0.7304 (1610) | ACCA=0.6961 (2155) |
| CACT=0.8175 (1694) | ACTA=0.8049 (509) | GACT=0.7297 (1132) | CGTA=0.6874 (552) |
| CGAG=0.8028 (2426) | AGCA=0.7988 (4445) | CGAA=0.7159 (2143) | TGTG=0.6844 (1849) |
| AATT=0.8019 (3416) | TCTA=0.7966 (342) | GAGT=0.7149 (1141) | GAAA=0.6757 (2726) |

**B**

| **Mutant-r3 type *D. melanogaster*** | | | |
|---|---|---|---|
| **Lowest 10 Pearson-correlation outliers and their motifs** | | | |
| R(10 bp) | R(50 bp) | R(100 bp) | R(200 bp) |
| GTAG=-0.0110 (4451) | GACT=-0.0155 (3709) | CCTA=-0.0079 (3680) | TACT=-0.0167 (3730) |
| TCTA=-0.0109 (4014) | ATAC=-0.0121 (3061) | ACTA=-0.0067 (3991) | TTAC=-0.0142 (3633) |
| CCTA=-0.0099 (3058) | CATA=-0.0108 (3062) | ACGT=-0.0028 (4259) | GGTA=-0.0133 (3456) |
| TAGG=-0.0090 (3009) | TAAC=-0.0100 (2946) | TCCC=-0.0013 (14111) | AGTG=-0.0114 (5061) |
| CCGA=-0.0085 (7526) | TAAG=-0.0086 (3286) | TCTA=-0.0010 (5028) | TCAC=-0.0113 (5532) |
| TAAC=-0.0067 (3901) | TCAT=-0.0073 (5180) | TAGA=-0.0007 (4971) | GTTA=-0.0104 (3718) |
| CGTA=-0.0057 (2955) | GCTA=-0.0069 (3236) | CGTA=0.0014 (3747) | CTCA=-0.0095 (5707) |
| GATA=-0.0045 (4236) | GATA=-0.0059 (3368) | ACGC=0.0019 (7405) | GTAA=-0.0093 (3437) |
| CTAC=-0.0034 (4526) | ATTG=-0.0058 (5479) | GATT=0.0027 (8208) | GTGT=-0.0092 (5366) |
| CTAG=-0.0025 (3236) | CCTA=-0.0058 (2559) | TAGT=0.0027 (3768) | AACC=-0.0084 (7436) |
| **Highest 10 Pearson-correlation outliers and their motifs** | | | |
| R(10 bp) | R(50 bp) | R(100 bp) | R(200 bp) |
| TAGT=0.5918 (3456) | TACC=0.2893 (2881) | TAAC=0.1998 (4402) | CGAA=0.1921 (7254) |
| AAGG=0.2979 (9354) | TAGC=0.2644 (3270) | CAAG=0.1738 (12860) | ACGG=0.1903 (5536) |
| CACT=0.2686 (5633) | CTTG=0.1926 (8830) | GCGT=0.1731 (7475) | CAAG=0.1680 (10024) |
| TATC=0.2670 (4379) | GTGT=0.1847 (4722) | TATC=0.1621 (5203) | TCAA=0.1667 (8631) |
| TGGA=0.2447 (13432) | TATG=0.1786 (3069) | AACA=0.1608 (14808) | CCCT=0.1633 (9198) |
| CAAG=0.2332 (10781) | CGGA=0.1735 (7791) | CCCG=0.1565 (14386) | AGGA=0.1460 (11653) |
| ATCC=0.2281 (8329) | CAAG=0.1729 (8732) | ATCG=0.1526 (7561) | CTTA=0.1453 (3943) |
| CCCT=0.2256 (9530) | GTTT=0.1697 (8514) | ACCA=0.1421 (13261) | CTTG=0.1422 (10212) |
| AATT=0.2210 (10274) | AATT=0.1612 (7287) | CTAG=0.1413 (4125) | GTAC=0.1415 (3391) |
| TACG=0.2165 (2974) | GGTA=0.1547 (2923) | CCAA=0.1409 (14643) | TACC=0.1394 (3586) |

Table S2: *D. melanogaster* Pearson correlation co-efficient outliers (top ten and lowest ten) for different *intra-exon 4-mer* motif sequence pairs at 10, 50, 100 and 200 bp spacings. A) 2nd Replicate from the wild-type dataset B) 2nd Replicate from the mutant-type dataset. Sample sizes are given in parenthesis.

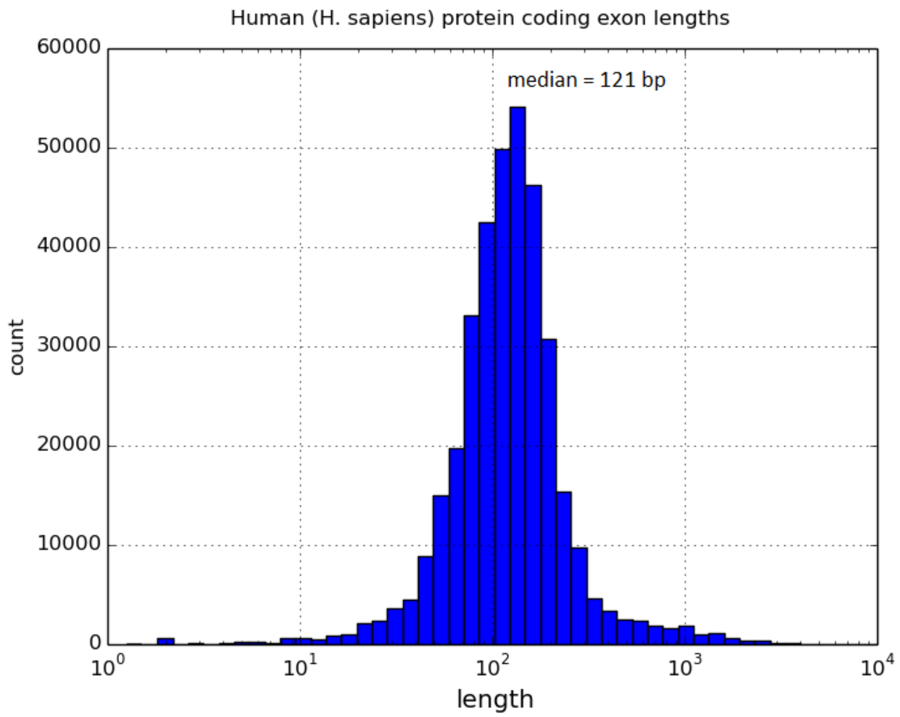


Figure S2: Histogram of *H. sapiens* Exon lengths . The data is computed from the hg19/GRCh37 genome annotation GTF which was used in the analysis of the IVT samples. The median exon length is 121 bp, mean 320 bp, Q1=84 bp, Q3=168 bp, the longest exon is 12218 bp.

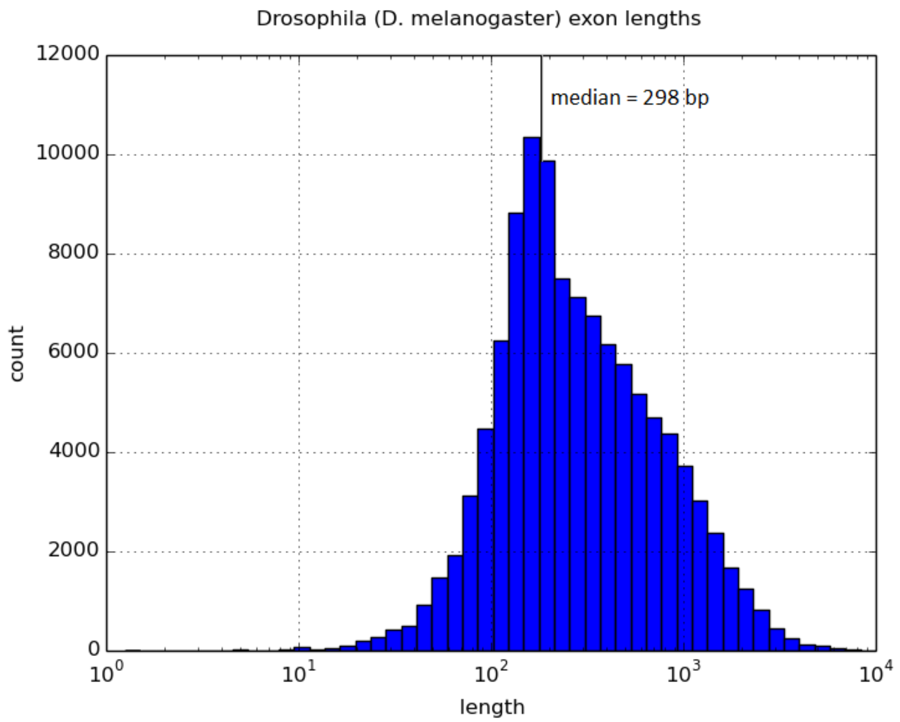


Figure S3: Histogram of Drosophila Exon lengths for both wild and mutant-r2 type *D. melanogaster*. The data is computed from the FlyBase genome annotation GTF (Version 6.15) which was used throughout the analysis. The median exon length is 298 bp, mean 540.1 bp, Q1=155 bp, Q3=637 bp, the longest exon is 28070 bp.

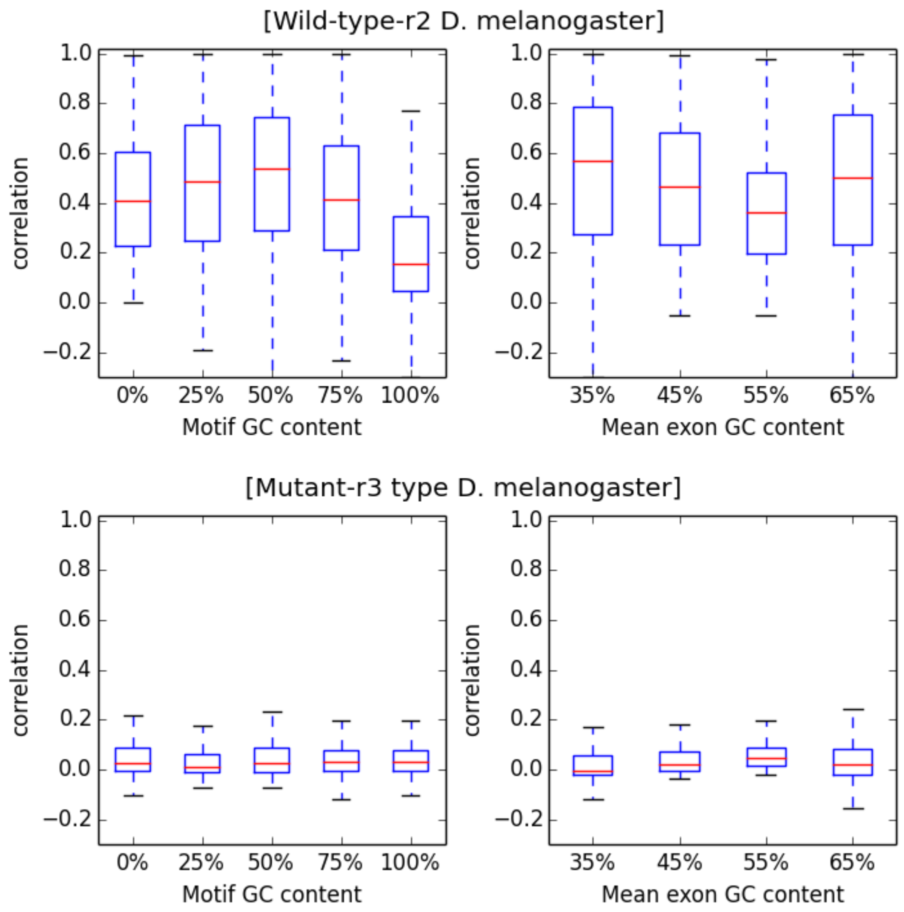


Figure S4: Correlation (Pearson's) as a function of *4-mer* motif and exon GC content for 2nd replicates of both wild and mutant-r2 *D. melanogaster* transcriptomes.
